# Dual inhibition of SHP2 and autophagy suppresses NF1-associated Malignant Peripheral Nerve Sheath Tumors

**DOI:** 10.1101/2022.08.18.504317

**Authors:** Sameer Farouk Sait, Kwan-ho Tang, Steve Angus, Rebecca Brown, Daochun Sun, Xuanhua Xie, Charlene Iltis, Michelle Lien, Nicholas Socci, Tejus Bale, Christopher Davis, Shelley A.H Dixon, Chi Zhang, D. Wade Clapp, Benjamin G. Neel, Luis F. Parada

## Abstract

Malignant peripheral nerve sheath tumors (MPNSTs) are aggressive sarcomas and the primary cause of mortality in patients with neurofibromatosis type 1 (NF1). MPNSTs develop within pre-existing benign plexiform neurofibromas (PNs). PNs are driven solely by biallelic *NF1* loss eliciting RAS pathway activation and respond favorably to MEK inhibitor therapy. Our analysis of genetically engineered and orthotopic patient-derived xenograft MPNST indicates that MEK inhibition has poor anti-tumor efficacy. By contrast, upstream inhibition of RAS through the protein-tyrosine phosphatase SHP2 reduced downstream signaling and suppressed *NF1* MPNST growth, although resistance eventually emerged. To investigate possible mechanisms of acquired resistance, kinomic analyses of resistant tumors was performed, and data analysis identified enrichment of activated autophagy pathway protein kinases. Combining pharmacological blockade of autophagy and SHP2 inhibition resulted in durable responses in *NF1* MPNSTs in both genetic and orthotopic xenograft mouse models. Our studies can be rapidly translated into a clinical trial to evaluate SHP2 inhibition in conjunction with autophagy inhibitors as a novel treatment approach for *NF1* MPNSTs.

**Statement of significance:** Currently, no effective therapies exist for MPNST. We demonstrate intrinsic MPNST resistance to MEKi monotherapy and identify SHP2 inhibition as an actionable vulnerability upstream of RAS. Furthermore, anti-tumor effects are extended and enhanced by dual exposure to autophagy pathway inhibition. Validation of these results as the most effective therapy to date in multiple genetically engineered models and in orthotopic patient-derived xenografts justify a clinical trial to evaluate SHP2i in conjunction with autophagy inhibitors as a novel treatment approach for *NF1* MPNSTs.

## Introduction

Neurofibromatosis type 1 (NF1) is an autosomal-dominant cancer predisposition syndrome with 50% of patients afflicted by multiple benign peripheral nerve sheath tumors termed plexiform neurofibromas (PNs) characterized by biallelic *NF1* loss (1). NF1 patients with PNs have an 8-13% lifetime risk of malignant transformation to an aggressive soft tissue sarcoma known as malignant peripheral nerve sheath tumor (MPNST) (2). MPNSTs harbor additional oncogenic events, that can include homozygous loss of *INK4A/ARF, P53* mutations, alterations in components of the PRC2 complex (*EED, SUZ12)*, or amplification of receptor tyrosine kinase (RTK) genes. The mortality caused by MPNSTs is high, approaching 100% in patients with unresectable, metastatic, or recurrent disease, as these tumors do not respond to current anti-cancer therapies (3).

The protein encoded by the *NF1* gene, neurofibromin, is a RAS-GTPase activating protein and homeostatic regulation of RAS protein by the Guanosine exchange factor/GTPase activating (GEF/GAP) cycle is disrupted in NF1 patients, resulting in elevated RAS-GTP levels and increased downstream RAF/MEK/ERK signaling. Preclinical studies of *Nf1*-mutant PN mouse models accurately predicted responses in clinical trials as exemplified by the success of single agent MEK inhibitors (4,5). By contrast, several monotherapy-based clinical trials in MPNST have yielded disappointing results, likely reflecting their increased genomic complexity (6).

The protein-tyrosine phosphatase SHP2 (encoded by *PTPN11*) functions as a positive signal transducer upstream of SOS1/2 and NF1 (7). SHP2 inhibition was recently demonstrated to inhibit RAF/MEK/ERK pathway activation and tumor growth in RAS-mutant preclinical models of solid tumors (8-11), and several SHP2 inhibitors (SHP2i) are currently in clinical trials either as monotherapy or in combination with other agents (12,13). Given its position upstream of SOS1/2, we hypothesized that SHP2 inhibitors could more effectively decrease RAS-GTP levels and downstream signaling in *NF1*-mutant tumors. We find that SHP2 inhibition can efficiently reduce RAS-GTP loading, block RAS-mediated RAF/MEK/ERK signaling and abrogate tumor growth in NF1-MPNST. Furthermore, SHP2 inhibition induces autophagic flux in resistant tumor cells, and pharmacological inhibition of autophagy enhances the effectiveness of SHP2 inhibition in mouse and human models of NF1-MPNST.

## Results

### SHP2 inhibition suppresses MPNST cell growth

To directly study the effects of Trametinib (MEK inhibitor, hereafter MEKi) versus SHP099 (SHP2 inhibitor, hereafter SHP2i) on MPNST, we established primary MPNST cultures from *Nf1-/-;Trp53-/- (NP) or Nf1-/-;Ink4a/Arf-/- (NI)* mutant mouse tumors (Supplementary Fig 1A) in defined, serum-free medium (14). Treatment with either inhibitor caused substantial *in vitro* cell growth arrest (Fig 1A and 1B). We next examined the biochemical effects and compared to baseline, MEKi-treated cells demonstrated increases in RAS-GTP, whereas SHP2i treatment resulted in decreased RAS-GTP at one hour and twenty-four hours (Fig 1C and 1D) (15,16). Further, MEKi resulted in a progressive increase in MEK1/2 Ser217/221 phosphorylation (p-MEK), while SHP2i reduced p-MEK levels (Fig 1E and 1F) (17). The initial reduction in phosphorylated ERK (p-ERK) levels following MEKi addition was partially restored by 48 hours in *NP* mutant MPNST, and to lesser extent, in *NI* mutant MPNST cultures (Fig 1E and 1F) (15-17). We also performed qRT-PCR analysis of multiple receptor tyrosine kinases (RTKs), which revealed selective upregulation of genes encoding several RTKs in MEKi versus SHP2i treated MPNST cells (Supplementary Fig 1B) (8). In aggregate, these data indicate that SHP2i effectively suppresses the Ras pathway (8,18).

**Figure 1.**
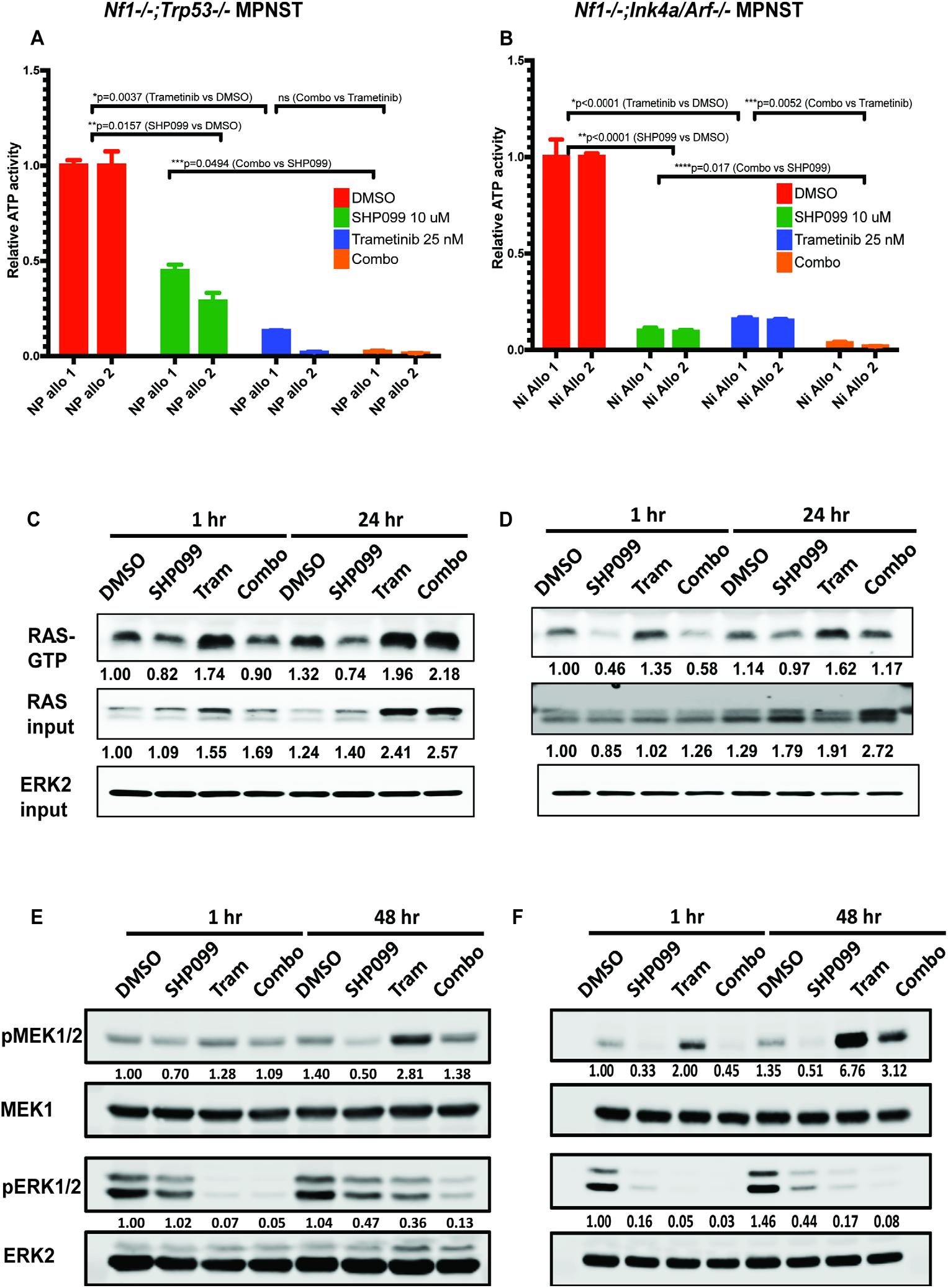
*In vitro* testing in mouse allograft models Cell titer glo viability assays of primary cultures of *NP* mutant **(A)** and *NI* mutant MPNST **(B)** (n=2 each) treated for 96 hr with vehicle (DMSO), Trametinib (25 nM), SHP099 (10 uM) or SHP099 and Trametinib. RAF-RBD pulldown assays performed after the indicated times of drug treatment in *NP* mutant **(C)** and *NI* mutant primary tumor cultures **(D)**. Immunoblots for the indicated analytes of lysates from *NP* mutant **(E)**; *and NI mutant* **(F)** primary cultures treated as indicated. Numbers under each lane represent quantification of p-MEK or p-ERK signals normalized to that of total MEK or ERK and compared to DMSO. Representative results from a minimum of three biological replicates are shown per condition.

### SHP2i exhibits enhanced mouse anti-tumor efficacy

We next turned to *in vivo* orthotopic genetically engineered MPNST models (GEMMs) derived from *NP* and *NI* genotypes to compare the anti-tumor effects of single agent MEKi (Trametinib 0.25 mg/kg daily; the human maximum tolerated dose equivalent allometrically scaled to mice) or SHP2i (SHP099 50 mg/kg daily), and dual MEKi/SHP2i (Trametinib 0.25 mg/kg daily, SHP099 50 mg/kg every other day) combination. The drug combination was toxic when full single agent doses were administered daily, as previously reported (8), requiring adjustments to the drug doses and schedules. Five thousand *NP* or *NI* MPNST cells each were injected orthotopically into twenty-four nude mice. Once tumors reached a volume of 250-500 mm^3^, mice were divided into four groups. Vehicle or drug was administered via oral gavage daily for a total of ten days, and tumor response was assessed. Waterfall plot illustrations showing the treated mouse tumor volumes relative to baseline at day ten demonstrated minimal anti-tumor effects for MEKi alone consistent with previous reports (17,19). In contrast, SHP2i monotherapy and the drug combination resulted in significantly reduced tumor volume (Fig 2A and 2B).

**Figure 2.**
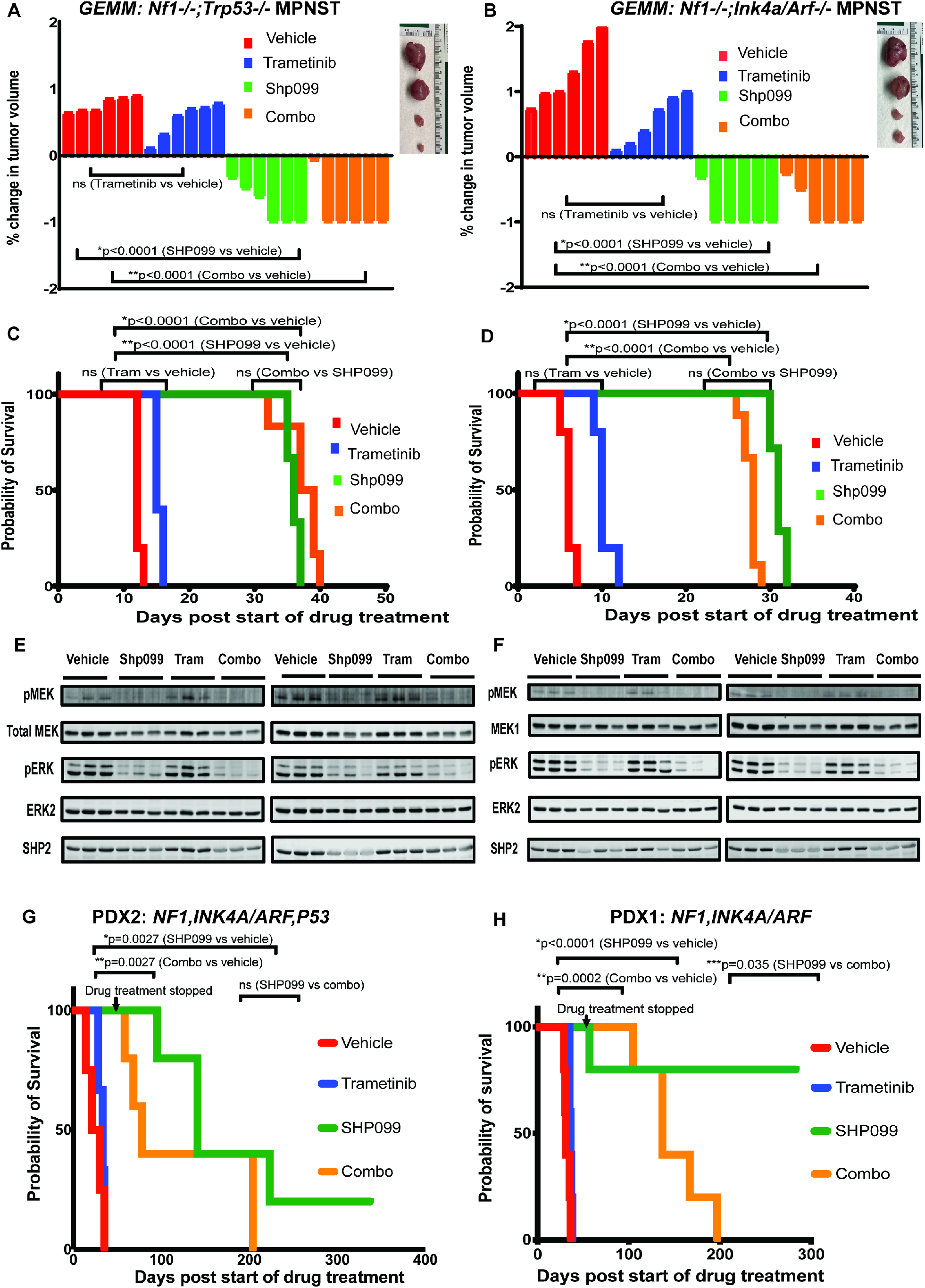
Effects of SHP2 or MEK inhibition alone or in combination in mouse MPNST GEMMs and patient-derived xenograft models Waterfall plots demonstrating change in tumor volume following 10 days of drug treatment relative to baseline in *NP mutant* **(A)** and *NI* mutant **(B)** MPNST models (n=6 mice per condition for each model) Kaplan-Meier curves showing survival of drug treated mice versus controls (vehicle) in *NP* mutant (**C)** and *NI* mutant **(D)** allograft models (n=6 mice per condition). Immunoblots of lysates obtained 6 hours after a single dose of drug treatment and 6 hours after three daily doses of drug treatment of mice bearing *NP* mutant **(E)** and *NI* mutant **(F)** MPNSTs (n=3 mice per condition). Numbers under each lane represent quantification of p-MEK or p-ERK signals normalized to total MEK or ERK signals and compared to DMSO. Kaplan Meier curve demonstrating survival analysis in drug treated mice versus controls (vehicle) in several independent NF1 MPNST PDX models (*PDX#1;* n=5 mice per condition) **(G)**, (*PDX#2*; n=3-5 mice per condition) **(H)**.

To assess the durability of anti-tumor effects, we next performed more extended experiments with drug treatment maintained until *NP* and *NI* MPNST reached a volume of 1000 mm^3^. Continuous therapy was well tolerated with no mice demonstrating obvious toxicity (data not shown). Consistent with the short-term assays above, SHP2i monotherapy and the drug combination provided significant delay in the time to tumor progression and extended survival by approximately threefold (Fig 2C and 2D). Immunoblot analyses were performed six hours following the first dose, and six hours following three consecutive daily doses, of vehicle, SHP2i, MEKi, and the combination. In both GEMMs, SHP2i monotherapy and combination therapy demonstrated effective suppression of p-ERK levels (Fig 2E and 2F). By contrast, MEKi monotherapy showed limited suppression of p-ERK levels in concordance with the minimal anti-tumor effects observed (Fig 2E and 2F). Analysis of additional RAS effector pathways indicated p-AKT suppression by SHP2i (Supplementary Fig 2A & 2B), and partial p-STAT3 suppression only in *NI* MPNST (Supplementary Fig 2B). The significance of these findings remains to be determined. Recent studies have highlighted variable p-AKT suppression by SHP2i across cancer cell lines (20). Consistent with the biological effects of SHP2i, immunohistochemical (IHC) analysis of *NP* and *NI* MPNST revealed decreased proliferation and induction of apoptosis as revealed by decreased Ki67 staining and increased cleaved caspase 3 (CC3) staining, respectively (Supplementary Fig 2C and Supplementary Fig 2D). SHP2i as monotherapy, or combined with MEKi, also suppressed tumor growth in a third spontaneous MPNST mouse model, which harbors mutations in the *Nf1* and *Trp53* genes (cis*NP)* (Supplementary Fig 2E) (14,21,22). Thus, in three independent GEMM models of NF1-MPNST, SHP2i monotherapy exhibits anti-tumor efficacy that significantly exceeds that of MEKi, as well as enhanced and extended suppression of the RAS/RAF/MEK/ERK pathway.

### SHP2i Suppression of Orthotopic *NF1*-MPNST PDX

To examine whether the SHP2i anti-tumor effects observed in MPNST GEMMs extend to the human disease, we generated NF1-mutant MPNST patient-derived xenografts (PDX) by orthotopic transplantation of surgical tumor specimens directly into the sciatic nerves (SN) of adult immunocompromised mice. Following the development of palpable tumors, the tumors were extracted, and fragments were used for re-transplantation (PDX-Passage 1), while additional fragments were used for pathological, immunohistochemical, biochemical, and genomic verification. Overall, our panel of orthotopic SN NF1-mutant MPNST PDX models (n=4) exhibit histological, molecular, and genomic fidelity to the index human tumors from which they were derived over three subsequent passages (Supplementary Fig 3A and 3B and data not shown).

We assessed the response of NF1-mutant PDX models to MEKi or SHP2i; and dual therapy at the same doses and schedules used for MPNST GEMMs. Treatment was initiated once tumors reached a volume of approximately 500 mm^3^, and drugs were delivered daily by oral gavage for twenty-one days, after which treatment was concluded, and tumors were evaluated. The waterfall plots represent the status of tumor volume relative to baseline at day twenty-one demonstrating the antitumor activity of SHP2i monotherapy (Supplementary Fig 4A and 4B). Consistent with these biological effects, immunohistochemical (IHC) analysis of PDX samples exhibited decreased proliferation (Ki67) and increased apoptosis (CC3; Supplementary Fig 4C) in SHP2i treated and combination-treated tumors compared to those receiving MEKi or vehicle.

To assess the durability of these anti-tumor effects, drug treatment was extended until tumors reached a volume of 1000 mm^3^ or until day sixty, whichever came first. While the drug combination delayed time to tumor progression, SHP2i monotherapy conferred a significant survival advantage across multiple independent NF1-PDX models (Fig 2G-H and Supplementary Fig 4D). Remarkably, these results included a few examples where SHP2i-treated PDX-bearing mice showed no evidence of tumors over time and were effectively cured of the disease (Fig 2G-H and Supplementary Fig 4D). Thus, consistent with the MPNST GEMMs, SHP2i has considerable anti-tumor properties in multiple genetically diverse patient-derived tumor models.

In summary, in the majority GEMMs and PDX models tested, SHP2i monotherapy provides superior MPNST suppression and survival in comparison to MEKi monotherapy or combination therapy. Nevertheless, with some exceptions, SHP2i resistance and tumor regrowth eventually emerges.

### *In vivo* resistance to SHP2i

To examine therapeutic resistance in greater detail, we turned to the less cumbersome MPNST GEMMs given their fidelity to the therapeutic responses seen in the PDX models. A typical response to SHP2i monotherapy in *NP* mutant MPNST is illustrated in Fig 3A. Upon SHP2i treatment, tumor volume initially regressed (weeks 1-2; *tumor shrinkage*) followed by a phase of *tumor stasis* (weeks 3-4) and eventually, tumor resurgence. We derived primary cultures from both SHP2i-resistant (*NP*-Res1 and *NP*-Res2) and SHP2i-sensitive vehicle (*NP-* V1 and *NP-*V2) treated tumors. The cell cultures showed similar growth rates in the absence of drug (Fig 3B). Surprisingly, the cells cultured from resistant tumors exhibited sensitivity to SHP2i (Fig 3B). However, when re-injected orthotopically, they formed tumors resistant to SHP2i. As expected, re-injected vehicle-treated MPNST cells gave rise to drug-sensitive tumors (Fig 3C). Similar results were seen with *NI* mutant MPNST allograft tumor cells with acquired *in vivo* resistance (*NI*-Res) to SHP2i and vehicle treated cells (*NI*-V) (Supplementary Fig 5A and data not shown). Thus, acquired resistance to SHP2i is observed only *in vivo* with native tumor stromal and microenvironmental components, and is not recapitulated in two-dimensional, defined primary culture conditions, wherein the cells continue to exhibit SHP2i sensitivity. Whether these differential *in vivo* and *in vitro* effects are cell autonomous, microenvironmental, or combinatorial, remains to be determined.

**Figure 3.**
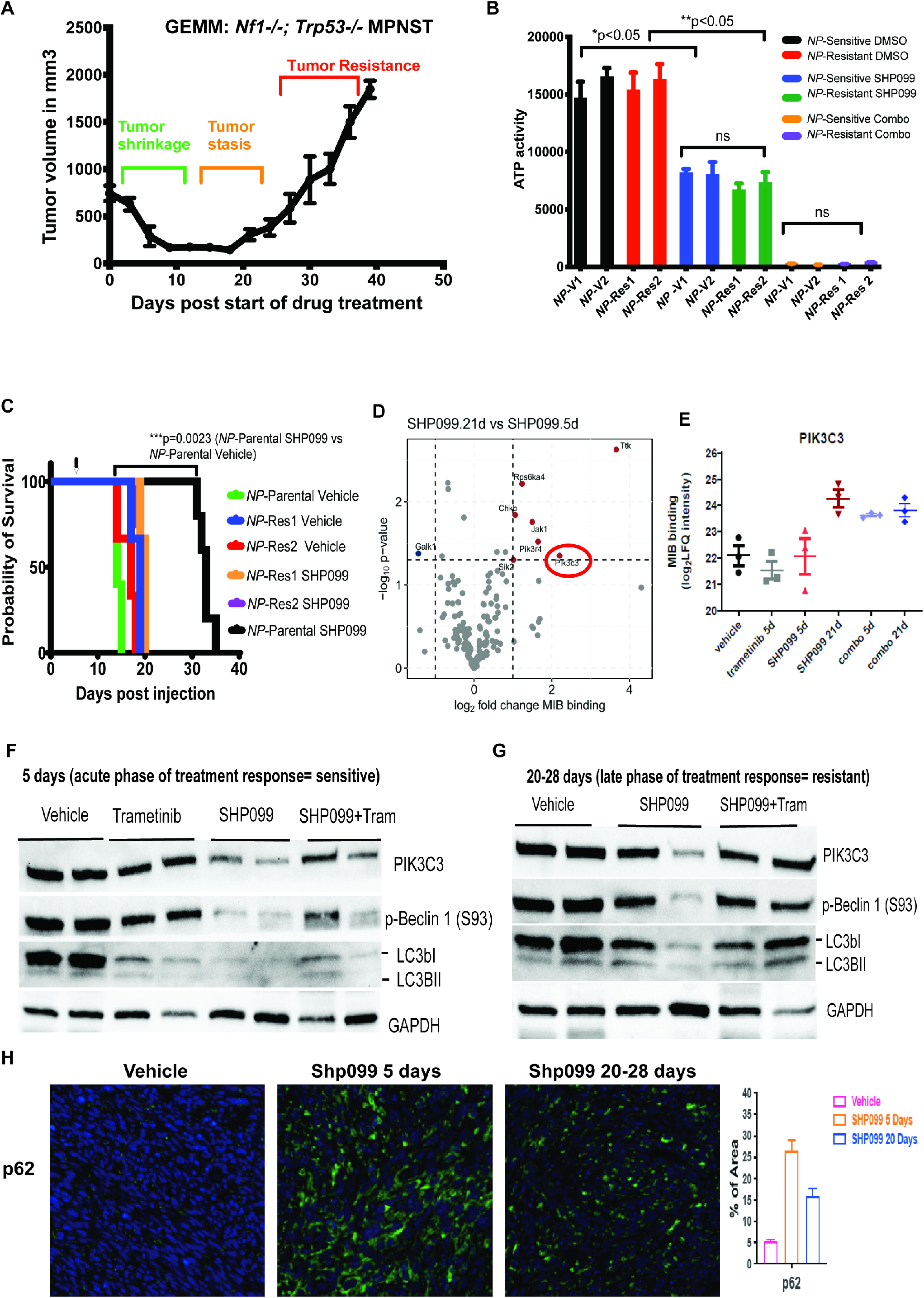
Resistance to SHP099 is associated with upregulation of the autophagy pathway *NP mutant* cells (*NP*-parental) were injected into the sciatic nerve of nude mice (n=5). When tumors reached 200-500 mm^3^, mice were treated daily with SHP099 (50 mg/kg) until resistance occurred, corresponding to approximately days 21-28 following start of therapy *(NP-*Res tumors). Tumor volumes in mm^3^ are presented in the graph (y axis) versus days of SHP099 treatment (x axis) (**A)**. Cell titer glo assays of *NP* mutant MPNST cells derived (*ex vivo*) from vehicle-treated (*NP*-V1 and *NP*-V2) or resistant (*NP*-SHP099Res1 and *NP*-SHP099Res2) tumors, exposed for 96 hours in vitro to DMSO, SHP099 (10 uM) or SHP099 (10 uM) and Trametinib (25 nM) **(B)** MPNST cells derived from a representative vehicle-treated (*NP*-V) or resistant tumor (*NP*-Res) were re-injected into the sciatic nerves of nude mice. SHP099 (indicated by arrow) or vehicle treatment was started at day 7 following cell injection as depicted by arrow (n = 3-5 mice per condition). Kaplan-Meier curves are shown **(C)**. Snap-frozen tumor from mice treated with vehicle, SHP099 (5 days=sensitive tumors), SHP099 (20-28 days=resistant tumors), Trametinib (5 days), combination of SHP099 and Trametinib (5 days=sensitive) and combination of SHP099 and Trametinib (20-28 days=resistant) were lysed and subjected to kinome profiling by multiplexed inhibitor bead/mass spectrometry (MIB/MS). Volcano plots show the log2 fold change in MIB binding (label-free quantification (LFQ) intensity determined by MS) plotted against the -log10 P-value for day 21 versus day 5 (resistant tumors compared to sensitive tumors). Horizontal dotted line, p=0.05 **(D)**. Log2 fold change in PIK3C3 binding to multiplex inhibitor beads in lysates from tumors treated with vehicle, Trametinib, SHP099 (5 days and 20-28 days) or combination of SHP099 and Trametinib (5 days and 20-28 days) **(E)**. Immunoblots of lysates from *NP* mutant MPNST tumors (n=2 mice per condition) treated with vehicle, SHP099, Trametinib, or combination of SHP099 and Trametinib for 5 days **(F)** or 20-28 days **(G)**. Numbers under each lane represent quantification of PIK3C3, p-BECLIN1 (S93), LC3-I, and LC3-II normalized to GAPDH and compared to DMSO. Immunofluorescence demonstrating p62 staining in sections from vehicle- or drug-treated tumors (mice bearing *NP* mutant MPNST following treatment with vehicle, SHP099 (5 days) or SHP099 (20-28 days) **(H)**. Panel at right shows quantification.

### SHP2i-resistant MPNSTs activate autophagy

The eventual acquisition of *in vivo* SHP2i resistance by MPNST could reflect specific transcriptional adaptations or alternatively, post-translational adaptations that might affect signaling networks. To examine potential clues at the post-translational level, we performed a kinomic analysis of fresh frozen tumor samples from vehicle-treated and drug-resistant MPNST. For each timepoint analyzed, we selected three independent tumor-bearing animals. Tumors from mice treated for 5 days with SHP2i represent early therapeutic responses (*tumor shrinkage*) (e.g., Fig 3A for *NP*-mutant MPNST allografts and data not shown for *NI* mutant MPNST allografts). To identify tumors in the earliest phases of therapy resistance (late *tumor stasis*), tumors were selected once palpable evidence of evolving therapeutic resistance and resumed tumor progression became apparent but prior to initiation of rampant tumor growth (20-28 days; e.g., Fig. 3A for *NP* MPNST and data not shown for *NI* MPNST). To identify candidate kinases associated with resistance to SHP2i (SHP099), we leveraged a chemical proteomic approach that utilizes multiplexed kinase inhibitor beads (MIBs), in which Sepharose beads are covalently coupled to broad-spectrum, Type I kinase inhibitors (23-26). MIB gravity-flow affinity chromatography, combined with mass spectrometry (MIB/MS), enables capture and identification of kinases in lysates on a kinome scale (23-26). Kinase binding to MIBs is influenced by their protein expression levels, by the affinity of a given kinase for the inhibitors used for MIB capture, and by their activity (26).

Snap-frozen tumor samples were subjected to MIB/MS kinome analysis, and the results are depicted as volcano plots. Notably, increased MIB binding of several kinases, including PIK3C3 (a.k.a., hVPS34), TTK, and LRRK2 was detected in SHP2i-treated *NP* and *NI* MPNSTs (Fig 3D & Supplementary Fig 5B). An obligate regulatory subunit of PIK3C3m PIK3R4 (a.k.a. hVPS15) also was recovered in the bead pulldowns (Fig 3D). Increased MIB binding of PIK3C3 kinase was especially pronounced in *NP* early resistant tumors (SHP2i at 20-28 days) compared with vehicle-treated control tumors or tumors treated with a short course of therapy (SHP2i at 5 days) (Fig 3E). hVPS34 (vacuolar protein sorting 34) is the only mammalian class III PI3K, and together with hVPS15 and BECLIN, catalyzes production of PI3 phosphate, a lipid important for autophagosome membrane formation (27).

To validate these observations, immunoblotting for autophagy pathway intermediates was performed. In both *NP* and *NI* MPNST models, initial treatment with SHP2i (i.e., *tumor shrinkage phase*) suppressed autophagic flux (AF), as evidenced by decreased levels of: phosphorylated BECLIN1 (Ser 93), PIK3C3, and LC3; and by increased p62 immunofluorescence staining (Fig 3F & 3H; data not shown and Supplementary Fig 5C) (28,29). Consistent with induction of autophagy, tumors treated for 21-28 days when early therapeutic resistance manifests (late *tumor stasis)*, showed elevated levels of p-BECLIN1 and LC3II, and decreased p62 staining (relative to 5 day treated tumors), (Fig 3G & 3H; data not shown and Supplementary Fig 5C) (28,29).

We also conducted RNA-seq experiments on the *NP* and *NI* MPNST mouse tumor samples treated with SHP2i for 5 days (early therapeutic response) and 20-28 days (evolved therapeutic resistance). We specifically focused on autophagy related genes, consisting of the essential genes involved in autophagy initiation, nucleation, and expansion (Supplementary Table 2) (30). Differential expression and pathway enrichment analysis demonstrated that consistent with the kinomic data, essential autophagy related genes are significantly downregulated in *NP* mutant MPNST tumor samples treated with SHP2i for 5 days (early therapeutic response) relative to vehicle treated tumors (Supplementary Fig 6A for *NP* mutant MPNST allografts and data not shown for *NI* mutant MPNST allografts). Conversely, essential autophagy related genes are significantly upregulated in the tumor samples treated with SHP2i for 20-28 days (evolved therapeutic resistance) relative to tumors treated with SHP2i for 5 days (early therapeutic response) (Supplementary Fig 6B for *NP* mutant MPNST allografts and data not shown for *NI* mutant MPNST allografts).

In summary, NF1 mutant MPNST exposed to SHP2i initially suppress tumor growth, autophagy, and undergo cleaved caspase 3 mediated apoptosis (Supplementary Fig 2B). However, with continued SHP2i, tumor growth resumes accompanied by increased autophagic flux.

### Pharmacologic inhibition of autophagy improves anti-tumor effectiveness

We investigated whether in MPNST, enhanced autophagy might be mechanistically linked to the emergence of SHP2i resistance. To test for an interplay, we therapeutically targeted autophagy using clinically available inhibitors. PIK3C3-IN1 is a highly potent, selective PIK3C3 inhibitor, whereas hydroxychloroquine (HQ) inhibits lysosomal acidification and blocks autophagy by affecting autophagosome fusion and degradation (31). To assess the combined effect of autophagy inhibition with SHP2i, primary MPNST cultures were treated with PIK3C3-IN1 (2.5uM) or HQ (25uM) in the presence or absence of SHP2i for four days and cell numbers were counted (see Methods for details). *NP* and *NI* MPNST cell number was reduced following combination treatment with SHP2i and HQ or PIK3C3-IN1 compared with single agent treatment (Supplementary Fig 7A-C).

We next assessed the effects of combination therapy in mice bearing *NP* MPNST. Nude mice were implanted with five thousand MPNST cells, divided into four cohorts, and treated with vehicle (control), HQ (50 mg/kg daily), SHP2i (SHP099 50 mg/kg daily), or the combination of SHP2i plus HQ (SHP099 50 mg/kg daily and HQ 50 mg/kg daily). All agents were administered by oral gavage once tumors had reached a volume of 500 mm^3^. As in the preceding experiments, SHP2i effectively inhibited tumor growth and delayed progression by several weeks. Single agent HQ had no detectable effect on tumor growth (32,33). However, the combination of HQ and SHP2i resulted in significant additional delay in tumor progression, without evidence of added toxicity (Fig 4A and data not shown).

**Figure 4.**
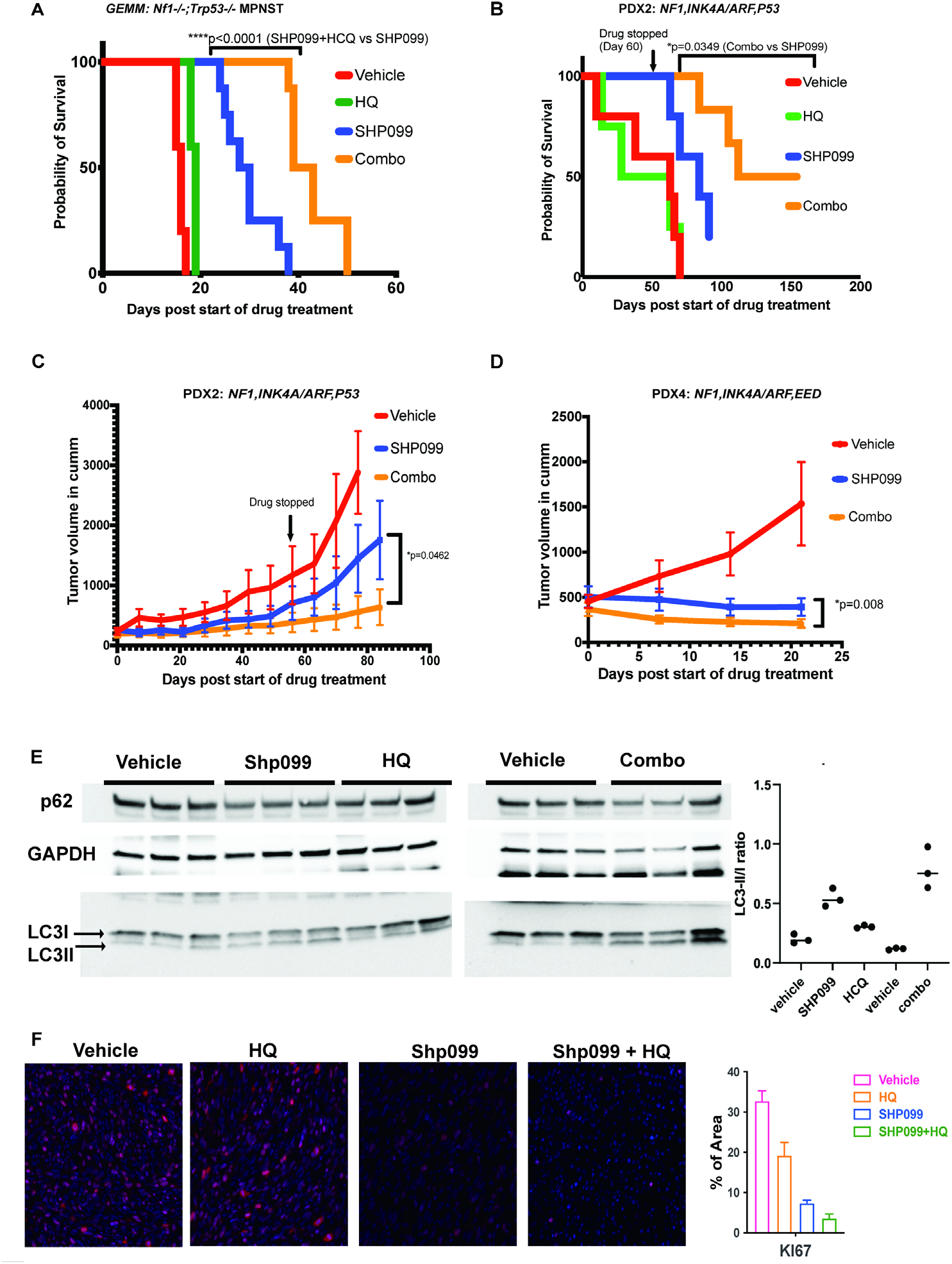
Effects of combined SHP2 and autophagy blockade in NF1 MPNST models Kaplan-Meier curves showing survival of drug treated mice versus controls (vehicle) in *NP* mutant MPNST (n=5-8 mice per group) **(A)** and an NF1 MPNST PDX model (PDX#*2*, n=4-6 mice group) **(B)**. Graphs depicting tumor volumes (Y axis) plotted against days of treatment with vehicle, SHP099 (50 mg/kg daily), or combination of SHP099 (50 mg/kg daily) and hydroxychloroquine (50 mg/kg daily) in mice implanted with NF1 MPNST PDX#2 tumors **(C)** (passage 1 xenografts; n=5-6 mice per condition) and mice implanted with PDX#4 tumors (passage 1 xenografts; n=5-6 mice per condition) **(D)**. Immunoblots of lysates obtained 6 hours after administration of a single dose of drug treatment of mice bearing *NP* mutant MPNSTs (n=3 mice per condition) **(E)**. Panel at right shows quantification of LC3II/I ratios. Immunofluorescence demonstrating Ki67 staining in tumor sections from mice bearing *NP* mutant MPNST and treated with vehicle, SHP099, HQ, or SHP099 and HQ (n=2 biological replicates) **(F)**. Panel at right shows quantification.

We also assessed the response of two genetically distinct NF1-MPNST PDX models to single agent SHP2i, HQ or the drug combination. Again, the drug combination demonstrated a significant survival advantage compared to SHP2i treatment alone (Fig 4B). Remarkably, three of six PDX#2 combination-treated mice had no detectable tumors at prolonged follow-up (160 days) despite discontinuation of therapy at day sixty (Fig 4B). Combined SHP2i and HQ treatment also resulted in deeper tumor regression (Fig 4C and 4D). Immunoblot analyses were performed six hours following a single dose of vehicle, SHP2i, HQ and the combination in mice injected with *NP* mutant MPNST. Combination treated tumors demonstrated increased LC3II levels (Fig 4E), increased levels of cleaved PARP indicative of cellular apoptosis (Supplementary Fig 7D) and decreased proliferation as demonstrated by lower Ki67 staining (Fig 4F). Snap-frozen tumor samples from mice implanted with PDX#4 tumors that received vehicle or drug treatment for 20 days were subjected to MIB/MS kinome analysis, and the results are depicted as volcano plots which demonstrate suppressed MIB binding of PIK3C3 kinase in drug combination treated tumors relative to SHP099 monotherapy treated tumors (Supplementary Fig 7E and 7F). Collectively, these data indicate that autophagy inhibition with HQ significantly improves the extent and durability of response to SHP2i both in GEMMs and diverse PDX models of NF1 mutant MPNST.

## Discussion

Genetically engineered mouse models of NF1 have exhibited remarkable fidelity in replicating the disease hallmarks (34,35). This has permitted otherwise impossible investigations into the origins of tumors, their interplay with the microenvironment, and more recently, prediction of clinical outcomes to PN therapies (4,26). NF1 mouse models have also contributed much insight into the origin, natural history, and genetic properties of MPNST, but no experimental therapies have yet been effective in clinical trials (3,36,37). We were prompted to reinvestigate RAS pathway inhibition for MPNST in the wake of recent preclinical and clinical success in the treatment of NF1 mutant PNs [58, 59]. The clinical impact of molecular therapies targeting the RAS/RAF/MEK/ERK pathway in other malignancies is highly variable dependent on the type of molecular alteration and/or the cellular and tissue context in which cancers arise (38). Certainly, in genomically complex and clinically aggressive diseases, combination with additional anti-cancer agents tends to increase efficacy (38). Given the well-established phenomenon of compensatory feedback upstream activation following RAS pathway inhibition (16,39-43), we reasoned that use of inhibitors upstream of RAS could perhaps mitigate such secondary activation in MPNST (26). Therefore, we re-examined MEKi in comparison with SHP2i in multiple NF1 MPNST mouse models. Only SHP2i monotherapy or combined therapy exhibited sustained RAS pathway inactivation and clear anti-tumor efficacy. We extended and validated these findings by using several recently developed orthotopic PDX models of NF1 MPNST with diverse mutational spectra. Several NF1 MPNST clinical trials are underway (NCT03433183) or planned (NCT05253131) which use a diverse array of combination therapies with MEKi as the backbone. Our results suggest that SHP2i (or nodes upstream of RAS such as SOS1/2) may provide a more effective backbone to test novel rational combinatorial treatments in lieu of MEKi.

As is the case following most anti-cancer therapies in the clinical setting, GEMM and PDX tumors frequently recurred following initial SHP2i-mediated regression (44,45). When therapy resistant MPNST tumors were placed in primary culture, they resumed SHP2i sensitivity yet when retransplanted orthotopically, the tumors remained SHP2i resistant and expanded aggressively. Thus, in MPNST, either cancer cell-intrinsic properties reconfigure to resume therapy resistance, or possible extrinsic/non autonomous factors including hypoxia, tumor stroma or microenvironment cells may exert unique anti-tumor influences such as the secretion of a growth factor that is not present in the culture medium and that bypasses SHP2i. With focus on adaptive intrinsic tumor cell responses, we examined transcriptional and proteomic kinome states. Concordant with transcriptional upregulation of autophagy pathway genes in early resistant tumors, the MIB/MS platform kinome analysis revealed differential activation of critical components of autophagy signaling. In the context of specific *BRAF*^*V600E*^ and RAS mutant tumor cell lines, it has been reported that use of RAF or MEK inhibitors elevated autophagy as an acquired resistance mechanism (46-49). In our study, co-targeting of SHP2 and the autophagy pathway enhances therapeutic efficacy *in vitro* and *in vivo*.

Our screening system for NF1-MPNST holds promise as a much-needed predictive preclinical platform. Tumor-derived primary cultures can be used to triage promising mono- or combination therapies followed by progression through orthotopic mouse tumors for rapid *in vitro* and *in vivo* screening, culminating in validation in orthotopic PDX models. As exemplified here, *in vitro* results do not always hold when extended to *in vivo* studies accentuating the need for progressive validation of results across the platforms. The use of mouse models in these studies further yielded critical insights into the tumoral mechanisms of drug resistance to allosteric SHP2 inhibition. The GEMMs provide an added advantage in future investigations into the potential role of the microenvironment in contributing to adaptive resistance to therapy. While here we show that tumor cell autonomous activation of autophagy is critical for the SHP2i resistant state, it is likely that external microenvironmental forces, including the immune and inflammatory systems, may also contribute and will require further investigation (50,51).

A recent study reported that SHP099 monotherapy was only modestly effective at halting MPNST growth, while genetic *PTPN11* (encoding SHP2) knockdown was more effective (17). Further, the combination of MEKi plus SHP2i had better outcomes in cell cultures. Several differences in our experimental approach could account for the differential outcomes. Wang *et al*. primarily employed long-established, high-passage MPNST cell lines maintained in RPMI or DMEM with supplements including fetal calf serum and presumably incubated in 20% O_2_ (17). Our cell experiments exclusively employed primary cultures directly derived from MPNST GEMMs or from patient resections, and maintained orthotopically in mouse sciatic nerves, a common location for spontaneous MPNST. Our primary cultures are maintained at low passage number in defined, serum-free medium with growth factor supplements and 5% O_2_. Multiple *in vivo* MPNST models demonstrated that combination of MEKi with reduced dose SHP2i on a continuous schedule was poorly tolerated and inferior to SHP2i alone both in sciatic nerve implanted GEMM tumors or PDXs. Taken in aggregate, we attribute the effects seen in combination therapy primarily to the action of SHP2i. The results from early phase clinical trials in RAS mutant tumors assessing safety and toxicity of combined SHP2i and MEKi which have overlapping toxicity profiles will require critical evaluation (NCT03989115) and underscore the importance of identifying the optimal combinatorial regimens in relevant preclinical model systems prior to initiating multicenter studies in a rare disease cohort such as *NF1* MPNST.

This study focused on cell autonomous MPNST responses to anti-RAS pathway therapy. The conclusion that RAS-upstream targets may have stronger and more durable anti-tumor effects warrants further investigation including delineation of the precise sources of feedback activation that are counteracted by SHP2i in these tumors. Presumably a subset of RTKs is one likely source. The fact that *in vivo*, SHP2i elicits autophagy leading to acquired resistance points to a promising therapeutic avenue in a space where none exist.

Our findings indicate that SHP2i has unprecedented suppression activity NF1-MPNST. Further, combined targeting of SHP2i with the autophagy pathway might offer better efficacy and less toxicity compared with drug combinations currently being tested to provide a first meaningful weapon against MPNST. These results merit evaluation in a prospective clinical trial.

## Methods

### Primary cell cultures and Reagents

Cells were cultured in serum-free medium supplemented with B27 and N2, plus EGF and FGF (10 ng/ml each) in 5% oxygen at 37C. Every culture was passaged for no longer than 4 weeks. Each primary cell culture was tested for mycoplasma contamination by PCR 5-7 days after each thawing. SHP099 (HY-100388A) was purchased from Med-ChemExpress, and trametinib (GSK1120212-S2673) was purchased from Selleckchem.

### Biochemical Assays

#### RAS Activity Measurements

RAS activity was assessed by GST-RBD pulldown, followed by immunoblotting with pan-RAS or RAS isoform–specific antibodies. Whole-cell lysates were resolved by SDS-PAGE, followed by transfer to Nylon membranes. Briefly, one ml of lysis buffer (25 mM Tris-HCl, pH 7.2, 150 mM NaCl, 5 mM MgCl_2_, 5% glycerol, 1% NP40) containing protease inhibitors was added to treated cells for 15 minutes on ice, and lysates were scraped from the plate and centrifuged at 14,000 rpm for 15 minutes at 4°C. Clarified lysates (0.25-3 mg) were added to pre-washed GST-tagged RBD glutathione agarose beads (30 μL) for 1h at 4°C under constant rocking. Beads were then centrifuged, washed once, and eluted in 30 μL of 2x SDS-PAGE sample buffer. Immunodetection of RAS proteins was carried out with pan-RAS (Ab-3; Calbiochem; 1:1,000) antibodies, with ERK-2 (sc-1647; Santa Cruz Biotechnology; 1:1000) used as a loading control.

### Immunoblotting

Immunoblots were performed with the indicated primary antibodies, followed by IRDye-conjugated secondary antibodies and visualization by LI-COR. Whole cell lysates were generated in modified radioimmunoprecipitation (RIPA) buffer (50mM Tris-HCl pH 8.0, 150mM NaCl, 2mM EDTA, 1% NP-40, and 0.1% SDS), supplemented with protease (40µg/ml PMSF, 2µg/ml antipain, 2µg/ml pepstatin A, 20µg/ml leupeptin, and 20µg/ml aprotinin) and phosphatase (10mM NaF, 1mM Na_3_VO_4_, 10mM β-glycerophosphate, and 10mM sodium pyrophosphate) inhibitors. Total lysate protein was resolved by standard SDS-PAGE and transferred in 1X transfer buffer and 15% methanol. Membranes were incubated with their respective primary and secondary antibodies labeled with IRDye (680nm/800nm) and visualized by using a LICOR. Immunoblots were performed from tumor tissue lysed as described below for MIB/MS kinome profiling. Equal amounts of total protein (by Bradford assay) were separated by SDS-PAGE, transferred to nitrocellulose membranes, probed with the indicated primary antibody and anti-rabbit or anti-mouse secondary antibodies coupled to HRP, detected by ECL, and imaged on a BioRad ChemiDoc and densitometry performed using BioRad ImageLab. Antibodies against p-MEK 1/2 (#9121; 1:1000), MEK1 (61B12; #2352; 1:1000), p-AKT (Thr308) (#4056; 1:1000), p-AKT (Ser473) (#9271; 1:1000), pan-AKT (#2920; 1:1000), and p-STAT3 (Tyr705) (#9145; 1:1000), p-Beclin-1 (Ser93) (D9A5G) rabbit mAb (#14717S), LC3B Antibody (#2775S), PI3 Kinase Class III (D4E2) Rabbit mAb (#3358T), Beclin-1 (D40C5) Rabbit mAb (#3495T), and SQSTM1/p62 (#5114) were obtained from Cell Signaling. Anti-SHP2 (sc-280; 1:1000), anti-GAPDH (sc-365062, 1:1000), and -ERK2 (sc-1647; 1:1000) antibodies were purchased from Santa Cruz Biotechnology. Anti-STAT1 antibody (#NB500-514) was purchased from Novus.

### IHC

Hematoxylin and eosin, p62, cleaved caspase-3 (Cell Signaling, D3E9), and Ki67 (Spring Biosciences, SP6) staining was performed on paraffin sections by the Experimental Pathology Shared Resource at Perlmutter Cancer Center (PCC).

### Quantitative RT-PCR

Total RNA was isolated by using the Qiagen RNeasy kit. cDNA was generated by using the SuperScript IV First Strand Synthesis System (Invitrogen). qRT-PCR was performed with Fast SYBR Green Master Mix (Applied Biosystems), following the manufacturer’s protocol, in 384-well format in C1000 Touch Thermal Cycler (Bio-Rad). Differential gene-expression analysis was performed with CFX Manager (Bio-Rad) and normalized to GAPDH expression. Quantification and statistics were performed in triplicate. Primers used are listed in Supplementary Table 1.

### Cell viability assays

MPNST cells (5000) from the *NP* and *NI* mutant models were cultured simultaneously in 96-well plates coated with fibronectin. When 50% cell confluency was noted in each well, DMSO (vehicle control), Trametinib (25nM), SHP099 (10 µM) or a combination of SHP099 (10uM) and Trametinib (25 nM) was added. Following 96 hours of drug treatment, the Cell Titer Glo (CTG) assay (Promega, catalog no. G7570) was utilized to assess cell viability. Statistical analysis between groups was performed using a two-tailed unpaired Student’s t-test.

### Cell counting

MPNST cells (5000) derived from the *NP* and *NI* mutant models were cultured in 96-well plates until 50% cell confluency as above. Then DMSO (vehicle control), HQ (25 uM), PIK3C3-IN1 (2.5 uM), SHP099 (10 uM), SHP099 (10uM) plus HQ (25 uM), or SHP099 (10 uM) plus PIK3C3-IN1 (2.5 uM) were added. Following 96 hours of treatment, cells were counted manually using a hemocytometer using trypan blue dye for exclusion to assess cell viability. Statistical analysis between groups was performed using a two-tailed unpaired Student’s t-test.

#### Animal Experiments

All mice were maintained under formal MSKCC Institutional Animal Care and Use Committee (IACUC) protocols (MSK IRB#15-06-007 and 15-06-009).

### NF1 GEMMs

Allograft models: We utilized a previously described mouse model harboring the *Nes CGD* transgene that permits conditional knockdown of tumor suppressors in neural crest-derived Schwann cell progenitors (SCPs) (14). These SCPs are enriched in in the boundary cap (BC) and the dorsal root ganglia (DRG) and represent the likely cell-of-origin population for *NF1* mutant neurofibromas and, by extension, MPNSTs (52). BC and DRG were harvested as described (14) from E13.5 embryos from *Nf1 f/f;Trp53 f/f*;*CGD (NP)* and *Nf1 f/f;INK4a/Arf f/f*;*CGD (NI)* mice. We chose these models because *NF1, P53*, and *INK4a/ARF* are the most frequently mutated cancer-related genes in human MPNSTs. Recombination was induced *in vitro* by 4-hydroxytamoxifen at 10 µM for 72 hours and confirmed by genotyping. BC & DRG cells were orthotopically injected into the sciatic nerves of nude mice to generate *BC&DRG;CGD* allograft models (Supplementary Fig 1A).

For some experiments, we employed another genetically engineered mouse model (GEMM) of MPNST (*CisNP* mouse MPNST), which develops tumors spontaneously upon loss of heterozygosity (LOH) at the compound heterozygous *Nf1*^*+/-*^ and *Trp53*^*+/-*^ loci (*cisNP*). (22,53,54)

#### GEMM tumor dissociation

Primary tumors were dissected and minced into small tissue blocks, and then dissociated by gentleMACS Octo Dissociator (Miltenyi Biotec) with 1g/ml collagenase (Sigma-Aldrich, Cat# C8051) with the preset protocol. The dissociated cells were washed in PBS 3 times and treated with blood cell lysate buffer (Sigma-Aldrich, Cat# R7757).

### NF1-MPNST xenografts

#### PDX generation and model validation

Patients or their designated legal guardian who consented to research collection of tissue (MSKCC IRB#12-245, NCT01775072) and biospecimen research studies (IRB#06-107) had tumor tissue collected for PDX implantation in accordance with IRB#15-281. Tumor tissue was processed by a member of the Precision Pathology Biobanking Center (PPBC) (55) and stored at room temperature in HypoThermosol FRS (Stem Cell Technologies) until implantation into the host mouse. For tumor implantation, NSG animals were anesthetized, and the flank was sterilized with 70% ethanol. Tumor was minced into small fragments (4-6 mm^3^), and only macroscopically viable tumor tissue was used for implantation in the right upper thigh (orthotopic implantation, OT). A skin incision over the right hind-leg cut was used to expose the gluteus superficialis and biceps femoris muscles. A small scissor was used to cut the connective tissue that connects the muscles, revealing a small cavity traversed by the sciatic nerve. A piece of tumor was fixed to the surface of the nerve using absorbable sutures taking care not to breach the epineurium. For the interior muscular layer pocket, simple interrupted absorbable sutures were used, and the skin incision was closed with sterile stainless-steel wound clips. Primary tumors were grafted orthotopically in at least three to five different mice in the first passage. Implanted mice were monitored closely for tumor engraftment and growth. When tumors reached a volume (TV) of ∼1000 mm^3^ or if signs of distress related to tumor were apparent, mice were euthanized and the tumor dissected, fragmented or dissociated, and iteratively implanted into additional mice for a maximum of 5 passages. Representative tumor fragments were obtained from the initial passage (P0) and subsequent passages (P1-5) and processed for sequence validation, viable cryobanking for future model re-establishment, and other downstream applications. NSG (Jackson Labs Strain # 005557) or NSG-H (Jackson Labs, Strain #026222) mice and all procedures described are performed under sterile, barrier conditions.

PDX tumor samples were sequenced using MSK-IMPACT, a hybridization capture-based next-generation sequencing clinical assay (56). In summary, this assay consists of the following standard workflow: reads are mapped using BWA MEM, indel-realigned and baseQ-recalibrated using GATK and then mutations are called using MuTect (v1.1.4) and SomaticIndelDetector (GATK v2.3.9). Copy number was computed by using the total copy number counts computed on bins of 100b and GC-normalized. Then the log tumor normal ratio was segmented using the Circular Binary Segmentation (CBS) method (57). Analysis code is available at: https://github.com/soccin/seqCNA

### *In vivo* studies

All *in vivo* studies were performed with both mouse model systems (*NP and NI* mutant MPNST) and PDXs. For GEMM experiments, *NI* and *NP* mutant MPNST cells (5000 cells) were injected into nude mice. For PDX experiments, tumor chunks of equal size were implanted adjacent to the sciatic nerve of nude mice. Mice were split into 4 groups once tumors had reached a volume of 250-500 mm^3^. Vehicle and drugs were administered by oral gavage at the following doses and schedules.

SHP099 and Trametinib combination studies: SHP099 50 mg/kg daily, trametinib 0.25 mg/kg daily or the combination (SHP099 50 mg/kg every other day and trametinib 0.25 mg/kg daily). SHP099 and HQ combination studies: Hydroxychloroquine (50 mg/kg daily), SHP099 (50mg/kg daily) or the combination of SHP099 (50 mg/kg daily) plus HQ (50 mg/kg daily).

#### Short term efficacy studies

Drugs were administered for a total of 10 days and mice were sacrificed. Tumor volume was measured by calipers and calculated according to the protocol from Jackson Laboratory http://tumor.informatics.jax.org/mtbwi/live/www/html/SOCHelp.html.

#### Survival analyses

Drug treatment was administered continuously (7 days/week) until day 60 or until mice were euthanized once tumors reached a volume of 1000 mm^3^.

Statistical analysis between groups was performed using a two-tailed unpaired Student’s t-test. Kaplan-Meier survival curves were analyzed using log-rank (Mantel-Cox) test in GraphPad Prism 9.

### Kinome analysis

GEMM: Nude mice injected with 5000 *NP* mutant MPNST cells were treated with vehicle, Trametinib (0.25 mg/kg daily) for 5 days, SHP099 (50 mg/kg daily) for 5 days (n=3) or until onset of resistance (21-28 days, n=3), combination of SHP099 and Trametinib for 5 days or until onset of resistance (21-28 days).

PDX: Nude mice injected with PDX#4 tumor chunks were treated with vehicle, SHP099 (50 mg/kg daily), HQ (50 mg/kg daily) or the combination of SHP099 and HQ for 20 days. Snap-frozen tumors were harvested from mice. Protein lysate was prepared and utilized for kinome profiling by MIB/MS (23-26).

### Multiplexed inhibitor bead chromatography and mass spectrometry

MIB affinity chromatography and MS were performed on snap-frozen tissue from tumor-bearing mice treated with vehicle, Trametinib, SHP099, HQ or drug combinations. Tissue was crushed by mortar and pestle in ice-cold MIB lysis buffer (50 mM HEPES, 150 mM NaCl, 0.5% Triton X-100, 1 mM EDTA, 1 mM EGTA, pH 7.5) supplemented with complete protease inhibitor cocktail (Roche) and 1% phosphatase inhibitor cocktails 2 and 3 (Sigma). Extracts were sonicated 3 × 10 s, clarified by centrifugation, and filtered with a syringe (0.22 µm) before quantifying protein concentration by the Bradford assay. Equal amounts of total protein were gravity-flowed over MIB columns in high-salt MIB lysis buffer (1 M NaCl). The MIB columns consisted of a 125-µl mixture of five type I kinase inhibitors: VI-16832, PP58, purvalanol B (58), UNC-00064-12, UNC00064-79, and BKM-120, which were custom synthesized with hydrocarbon linkers and covalently linked to ECH-Sepharose (23-26). Bound protein was eluted twice with 0.5% SDS, 1% β-mercaptoethanol, 100 mM Tris-HCl, pH 6.8 for 15 min at 100C, and then treated with dithiothreitol (5 mM) for 25 min at 60°C and 20 mM iodoacetamide for 30 min in the dark. Following concentration by centrifuging with Amicon Ultra-4 filters (10-kDa cutoff) to ∼100 µl, samples were precipitated with methanol–chloroform, dried in a SpeedVac, and resuspended in 50 mM HEPES (pH 8.0). Tryptic digests were performed overnight at 37°C, extracted four times with 1 ml ethyl acetate to remove detergent, and dried, and peptides were passed through C18 spin columns according to the manufacturer’s protocol (Pierce). Peptides were resuspended in 0.1% formic acid and equal volume (approximately 20-0% of the final peptide suspension for each experimental set) was injected onto a Thermo EASY-Spray 75 μm × 25 cm C18 column and separated on a 120-min gradient (2–40% acetonitrile) using an EASY-nLC 1200 instrument. The Thermo Orbitrap Exploris 480 MS ESI parameters were as follows: Full Scan – Resolution 120,000, Scan range 375-1500 m/z, RF Lens 40%, AGC Target - Custom (Normalized AGC Target, 300%), 60 sec maximum injection time, Filters – Monoisotopic peak determination: Peptide, Intensity Threshold 5.0e3, Charge State 2-5, Dynamic Exclusion 30 sec, Data Dependent Mode – 20 Dependent scans, ddMS^2^ – Isolation Window 2 m/z, HCD Collision Energy 30%, Resolution 30,000, AGC Target – Custom (100%), maximum injection time 60 sec. Raw files were processed for label-free quantification (LFQ) by MaxQuant (version 1.6.11.0) LFQ using the appropriate UniProt/Swiss-Prot human or mouse database with fixed carbidomethyl (C) and variable oxidation (M) and acetyl (protein N-terminal) modifications, match time window 3 min. LFQ intensities for all annotated kinases with at least two razor+unique peptides were imported into Perseus software (version 1.6.10.50), log_2_ transformed, filtered to include annotated kinases with at least three valid values in one treatment group, and missing values imputed from the normal distribution for each column using default parameters (width 0.3, down shift 1.8) Two sample, unpaired Student t-tests of log_2_LFQ intensities were performed and plotted using R (*P*<0.05 significance cut-off). The mass spectrometry proteomics data have been deposited to the ProteomeXchange Consortium via the PRIDE (59) partner repository with the dataset identifier PXD035998.

Reviewer account details:

Username

Password: IoLPeTN6

### Immunostaining

Primary tumors were fixed overnight in 4% paraformaldehyde (PFA), then processed and embedded in paraffin. Sections (5 µm) were cut by using a microtome, stained with H & E, or subjected to immunofluorescence or horseradish peroxidase-based immunohistochemistry, as described (60). IF and IHC staining was quantified with ImageJ. Staining quantification was assumed to be normally distributed, but this was not formally tested.

### RNA Isolation and Sequencing

RNA was extracted from mice with *NP* mutant MPNST tumors treated with SHP2i for 5 days or 20-28 days (n=3 animals each). A 20-30 mg portion of *NP* mutant MPNST flash-frozen tumor tissue was used for RNA extraction using the Qiagen RNeasy Plus kit essentially as described by the manufacturer’s protocol. Tissue homogenization/disruption was by mortar and pestle and Buffer RLT Plus supplemented with beta-mercaptoethanol followed by passing homogenized samples over Qiagen QIAshredder columns. Qiagen RNase-Free DNase Set was used for on-column DNA digestion. Total RNA was submitted to the IU Center for Medical Genomics (CMG) for library preparation and sequencing. Total RNA was first evaluated for its quantity, and quality, using Agilent Bioanalyzer 2100. For RNA quality, a RIN number of above 8 was achieved. One hundred nanograms of total RNA was used. cDNA library preparation included mRNA purification/enrichment, RNA fragmentation, cDNA synthesis, ligation of index adaptors, and amplification, following the KAPA mRNA Hyper Prep Kit Technical Data Sheet, KR1352 – v4.17 (Roche Corporate). Each resulting indexed library was quantified and its quality accessed by Qubit and Agilent Bioanalyzer, and multiple libraries pooled in equal molarity. The pooled libraries were then denatured, and neutralized, before loading to NovaSeq 6000 sequencer at 300pM final concentration for 100bp paired-end sequencing (Illumina, Inc.). Approximately 30-40M reads per library were generated. A Phred quality score (Q score) was used to measure the quality of sequencing. More than 90% of the sequencing reads reached Q30 (99.9% base call accuracy).

### RNA-seq processing and analysis

The sequencing data was next assessed using FastQC (Babraham Bioinfomatics, Cambridge, UK) and then mapped to the mouse genome (UCSC mm10) using STAR RNA-seq aligner with the parameter: “—outSAMmapqUNIQUE 60”. Uniquely mapped sequencing reads were assigned to mm10 refGene genes using featureCounts (from subread). Genes with read count per million (CPM) >0.5 in more than 3 of the samples were kept. The data were further normalized using RPKM (Reads Per Kilobase Million). Differential gene expression analysis was conducted by using DESeq2 method with an FDR less than 0.05 as the significant cutoff (61). Pathway enrichment analyses were conducted by hypergeometric tests against mouse Gene Ontology and selected autophagy genes (Supplementary table 2), with P value less than 0.01 as the significant cutoff (62). The raw data have been submitted to the NCBI GEO database (accession number XXXX).

### Statistical Analysis

Data are expressed as mean ± standard deviation. Statistical significance was determined using Student t test, Mann–Whitney U test or one-way ANOVA, as appropriate. Statistical analyses were performed in Prism 9 (GraphPad Software). Significance was set at P = 0.05.

## Acknowledgments

The authors would like to thank members of the Parada laboratory for helpful suggestions and discussion. We thank the MSKCC Molecular Cytology Core and the DHART SPORE OMICS CORE for their assistance. The authors would like to acknowledge Alicia Maria Pedraza, Samhita Bapat, and Yanjiao Li for their contributions to this project. Immunohistochemistry was performed in part at the Experimental Pathology Shared Resource at Perlmutter Cancer Center supported by 3P30CA016087-41. S.F.S. is a recipient of the NCI SPORE U54 CA196519-01 Career Development Award and K12 Paul Calabrese Career Development Award in Clinical Oncology and L.F.P. is a recipient of Investigator-Initiated Research Award, Congressionally Directed Medical Research Programs from Department of Defense (W81XWH-16-1-0186); NCI SPORE U54 CA196519-01; R01: CA131313; NIH/NCI Cancer Center Support Grant P30 CA008748 and holds the Albert C. Foster Chair in Cancer Research. BGN is a recipient of CA49152.

